# Non-parametric synergy modeling with Gaussian processes

**DOI:** 10.1101/2021.04.02.438180

**Authors:** Yuliya Shapovalova, Tom Heskes, Tjeerd Dijkstra

**Affiliations:** Institute for Computing and Information Sciences, Radboud University, Postbus 9010, 6500 GL, Nijmegen, The Netherlands; Max Planck Institute for Developmental Biology, Max-Planck-Ring 25, D-72076, Tübingen, Germany; Dept. for Women’s Health, University Clinic Tübingen, Calwerstrasse 7, D-72076, Tübingen, Germany

## Abstract

**Background:** Understanding the synergetic and antagonistic effects of combinations of drugs and toxins is vital for many applications, including treatment of multifactorial diseases and ecotoxicological monitoring. Synergy is usually assessed by comparing the response of drug combinations to a predicted non-interactive response from reference (null) models. Possible choices of null models are Loewe additivity, Bliss independence and the recently rediscovered Hand model. A different approach is taken by the MuSyC model, which directly fits a generalization of the Hill model to the data. All of these models, however, fit the dose-response relationship with a parametric model.

**Results:** We propose the Hand-GP model, a non-parametric model based on the combination of the Hand model with Gaussian processes. We introduce a new logarithmic squared exponential kernel for the Gaussian process which captures the logarithmic dependence of response on dose. From the monotherapeutic response and the Hand principle, we construct a null reference response and synergy is assessed from the difference between this null reference and the Gaussian process fitted response. We evaluated performance of our model on a simulated data from Greco, two simulated data sets of our own design and two benchmark data sets from Chou and Talalay. We compare the Hand-GP model to standard synergy models and show that our model performs better than these standards. We also compare our model to the MuSyC model as example of a recent method which also fits a complete dose-response surface. Also in this case, the Hand-GP model performs better.

**Conclusion:** The Hand-GP model is a flexible model to capture synergy. Its non-parametric natures allows it to model a wide variety of response patterns.

## Background

Assessing synergy and antagonism of chemical compounds has applications in medicine, pharmacology and ecotoxicology. By comparing expected non-interactive (null) response and observed response, one can assess whether there is synergy or antagonism between two drugs. The most common candidates for the non-interactive response models are the Loewe additivity and Bliss independence models. However, there are other candidates such as the Highest Single Agent (HSA), the Tallarida and the recently rediscovered Hand model [20].

Loewe additivity and Bliss independence models often serve as a base for various extensions that incorporate more complex interaction patterns. [13] developed models for testing level-dependent and ratio-dependent synergy/antagonism. Level-dependent synergy/antagonism occurs when the difference (between non-interactive response and observed response) at low doses deviates from the difference at high doses. For example, antagonism can be observed at low doses and synergy and high doses. Ratio-dependent synergy/antagonism happens when, say, antagonism is observed when the mixture is dominated by drug 1 and synergy when the mixture is dominated by drug 2. [21] study asymmetric interactions of drugs. In particular, they define perpetrator and victim drugs. Perpetrators cause the change of *EC*_50_ of the other drug in the mixture, and victims are affected by these change. Both [13] and [21] develop methods for both Loewe additivity and Bliss independence type of models.

Most null reference models, Loewe additivity and Bliss independence in particular, are based on monotherapeutic dose-response curves. Frequently the Hill curve is chosen for modeling the monotherapeutic dose-response relationship [11]. Some studies have shown that other choices of monotherapeutic dose-response curves might be preferable in some cases [9, 14], but the Hill curve is the most common. Importantly, all these model are parametric meaning that they specify a fixed set of possible shapes as defined by the range of the parameters. Parametric models have advantages in that they are generally more interpretable than non-parametric models and perform well when the data follow the pattern implied by the parametric model [9, 14]. However, parametric models also have well-documented disadvantages, the most important one being their fixed set of possible shapes, when data behave differently from the parametric model assumptions. Hence, non-parametric models and especially Gaussian process models have become popular recently. For example, even in cases where a good but high-dimensional model is available from physics or engineering, GP’s have found applications as the workhorse of surrogate modeling [7]. Thus, especially for complicated systems like a biological cell’s response to a perturbation, the GP framework appears to be a natural approach.

As main competitor to our Hand-GP model, we use the recent MuSyC model [17] as this (1) also fits the entire response surface and (2) is highly parametric, with 12 parameters to specify the full model. The advantage of this model is that the parameters are interpretable and can be related to the hypothetical underlying mass-action rate equations [17]. Of the 12 parameters, 5 relate to synergy. Thus, a second advantage is that complicated synergy patterns can be captured in the parameters, say antagonism in efficacy (the effect for high doses) and synergy in potency (the 50% effect dose). For comparison, our Hand-GP model has only 4 hyperparameters, one of which captures the noise level, i.e. the lack of fit. We follow common machine learning terminology where the parameters of the GP are called hyperparameters, as they can be interpreted as such in a Bayesian setting.

Our proposed model is based on Gaussian processes (GP) and is non-parametric. A Gaussian process is completely defined by its mean and kernel functions. Different kernels can be used to express different structures observed in the data [19]. We propose a new kernel ideally suited to capture the logarithmic dependence of effect on compound dose in biochemical systems. As an extra benefit, the length scale hyperparameters of this kernel allows for data-adapted plotting of response curves and surfaces striking a middle ground between linear and logarithmic axes. In contrast to standard approaches to synergy, we fit a GP to all data instead of fitting only the monotherapeutic data, making the estimated monotherapeutic response more robust. We construct the null reference model numerically using the Hand model [20] by locally inverting the GP-fitted monotherapeutic data. Synergy is then assessed by a synergy effect surface as the difference between the GP-fitted response surface and the Hand-constructed null reference surface. This synergy effect surface allows in principle for different effects at different locations, for example dose-dependent synergy.

## Results

We compare the performance of the Hand-GP model with the MuSyC model. We also provide results of the original analysis with Loewe, Bliss, and Median Effect models when relevant. We analyse the performance of the models on three simulated and two experimental data sets. In detail, we use a simulated data set from [8] to which we refer as Greco data, two data sets from our own hand (one with strong synergy and one with strong antagonism), and two experimental data sets used by [4] to showcase their Median Effect model to which we refer as Chou and Talalay data.

All the data sets are inhibitory, meaning a larger compound dose leads to a smaller response. In the Hand-GP model, we quantify synergy by taking the difference between a response surface (GP fitted to the raw response data) and a null reference model, generated by the Hand construction from the monotherapeutic GP fitted response data. As generally speaking synergy is a desired effect, we subtract the GP-fitted response data from the null reference. Then, a positive difference means a smaller response than expected from the null reference, or equivalently a larger effect, i.e. more inhibition.

The MuSyC model was chosen as the main competitor of the Hand-GP model. We fit the MuSyC model using the Python library called synergy [22]. In the MuSyc model the response is fitted with a 12-parameter model. Synergy is then determined from a subset of the parameters, termed *β*, *α*_12_, *α*_21_, *γ*_12_ and *γ*_21_. Parameter *β* corresponds to change in synergistic efficacy, i.e. at large doses of both drugs the effect is *β* larger. Parameters *α* correspond to a change of the effective dose and *γ* to a change in the Hill slope coefficient. To enable a visual comparison of the Hand-GP model with the MuSyC model, we also fit a MySyc Model with *α*_12_ = *α*_21_ = *γ*_12_ = *γ*_21_ = 1 and *β* = 0. This constrained MuSyC model serves as an equivalent null reference model for direct visual comparison to our Hand-GP model. Subtracting the 12-parameter MuSyC model from the constrained MuSyC model, we obtain an effect surface for comparison to the effect surface of the Hand-GP model.

One of our contributions is the logarithmic squared exponential kernel, tailored for dose-response modeling. In Figure 1 we compare fits with our new kernel to fits with the standard (linear) squared exponential kernel. The data in this figure come from [8] and are discussed in more detail in the next section. The mean squared errors (MSE’s) of the fits with the logarithmic squared exponential kernel (72.01 for drug 1 and 63.35 for drug 2) are considerable lower than those of the fits with the (linear) squared exponential kernel (91.52 for drug 1 and 72.15 for drug 2). This shows that a Gaussian process with the logarithmic squared exponential kernel can approximate these data better.

**Figure 1.**
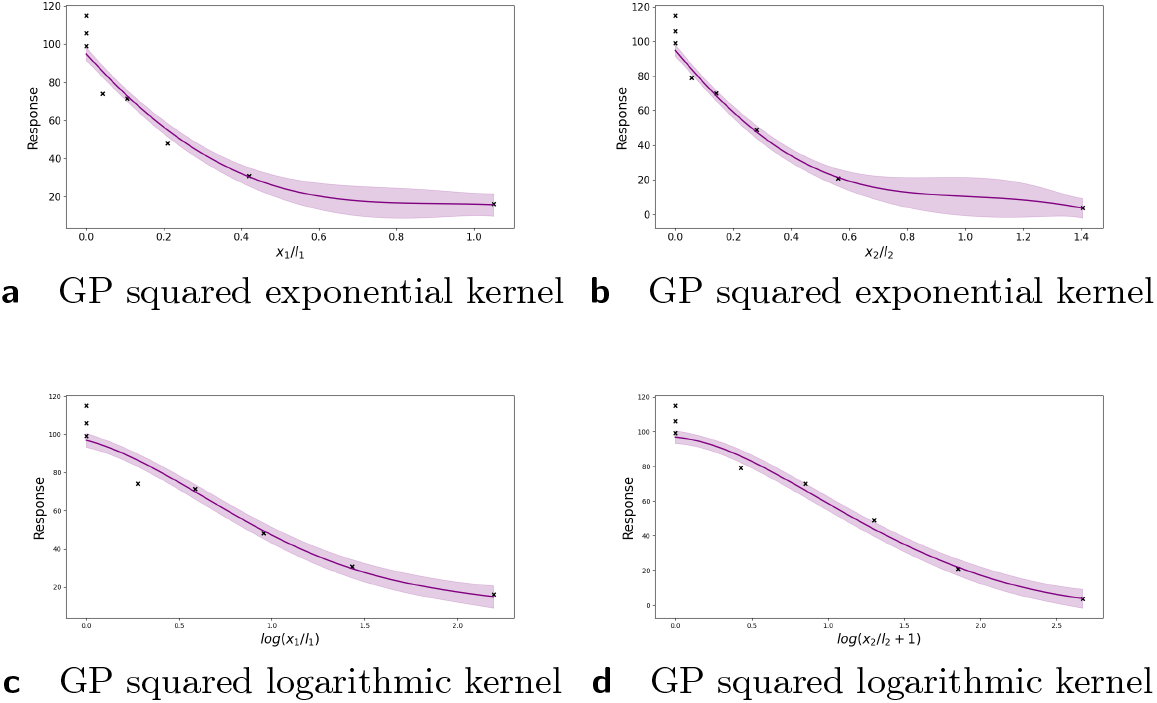
Dose-response curves for the Greco data using squared exponential and logarithmic squared exponential kernels. For both kernels we fit a complete dose-response surface, here only monotherapeutic slices are plotted. Left side: results for drug 1, right side: results for drug 2. Results from both Gaussian processes regressions are plotted on their natural scale: as (linear) *x*_*i*_/*l* for the squared exponential kernel and log(*x*_*i*_/*l*_*i*_ + 1) for the logarithmic squared exponential kernel. Note the difference in the estimates of the uncertainty.

### Greco simulated data

In this section, we illustrate the performance of the model on the benchmark data from Table 3 of [8]. The data set was generated with mild Loewe synergism (synergy coefficient 0.5), which in many regions of the response surface corresponds to mild Bliss antagonism. [8] considered 13 different models and reported detailed results for Loewe additivity and Bliss independence models which we compare to the Hand-GP and MuSyC models in Table 1. The Hand-GP model (third column of Table 1) predicts synergy except for a single dose combination (*x*_1_ = 5, *x*_2_ = 5) where it predicts antagonism. The MuSyC model (fourth column of Table 1) predicts antagonism for 15 dose combinations and synergy for 10. The Hand-GP model is clearly superior. The Hand-GP model performs even better than the Loewe model (fifth column of Table 1) although the data were simulated from this model, as the Loewe model incorrectly predicts antagonism for four dose combinations. The last (sixth) column of Table 1 shows results from Bliss independence which predicts antagonism except for three dose combinations. As the data were simulated from a Loewe model these results are not surprising and are discussed in detail by [8]. We present parameter estimates and overall effect estimates for both Hand-GP and MuSyC model in Table 2. As the Hand-GP model is non-parametric, we created an overall effect measure through the difference of the volumes under the surfaces of the unconstrained GP model and Hand-GP model. This effect measure is presented in the upper block and last column of Table 2 and indicates overall synergy. In this table, as well as in later tables, we color-code synergism as green, antagonism as red and additivity (no interaction effect) as grey. The MuSyC model has five parameters that correspond to different types of synergism/antagonism, thus each of them is color-coded separately. A MuSyC model that corresponds to no interaction, so just an additive effect, has parameters *β* = 0 and *α*_12_ = *α*_21_ = *γ*_12_ = *γ*_21_ = 1. We can see from Table 2 that additivity is predicted by all parameters of the MuSyC model as the additive effect values are within the 95% confidence intervals of each of these five parameters.

**Table 1.** Comparison of Hand-GP model to MuSyC model and Loewe and Bliss models from Greco [8] on the Greco simulated data

| $x_1$ | $x_2$ | Hand-GP | MuSyC | Loewe | Bliss |
| --- | --- | --- | --- | --- | --- |
| 2.0 | 0.2 | SYN | ANT | ANT | ANT |
| 2.0 | 0.5 | SYN | SYN | SYN | ANT |
| 2.0 | 1.0 | SYN | ANT | ANT | ANT |
| 2.0 | 2.0 | SYN | SYN | SYN | ANT |
| 2.0 | 5.0 | SYN | ANT | SYN | ANT |
| 5.0 | 0.2 | SYN | ANT | SYN | ANT |
| 5.0 | 0.5 | SYN | ANT | SYN | ANT |
| 5.0 | 1.0 | SYN | SYN | SYN | SYN |
| 5.0 | 2.0 | SYN | ANT | ANT | ANT |
| 5.0 | 5.0 | ANT | ANT | SYN | ANT |
| 10.0 | 0.2 | SYN | ANT | ANT | ANT |
| 10.0 | 0.5 | SYN | ANT | SYN | ANT |
| 10.0 | 1.0 | SYN | SYN | SYN | SYN |
| 10.0 | 2.0 | SYN | SYN | SYN | ANT |
| 10.0 | 5.0 | SYN | ANT | SYN | ANT |
| 20.0 | 0.2 | SYN | SYN | SYN | ANT |
| 20.0 | 0.5 | SYN | SYN | SYN | ANT |
| 20.0 | 1.0 | SYN | ANT | SYN | ANT |
| 20.0 | 2.0 | SYN | ANT | SYN | ANT |
| 20.0 | 5.0 | SYN | ANT | SYN | ANT |
| 50.0 | 0.2 | SYN | ANT | SYN | ANT |
| 50.0 | 0.5 | SYN | ANT | SYN | ANT |
| 50.0 | 1.0 | SYN | SYN | SYN | SYN |
| 50.0 | 2.0 | SYN | SYN | SYN | ANT |
| 50.0 | 5.0 | SYN | SYN | SYN | ANT |

**Table 2.**
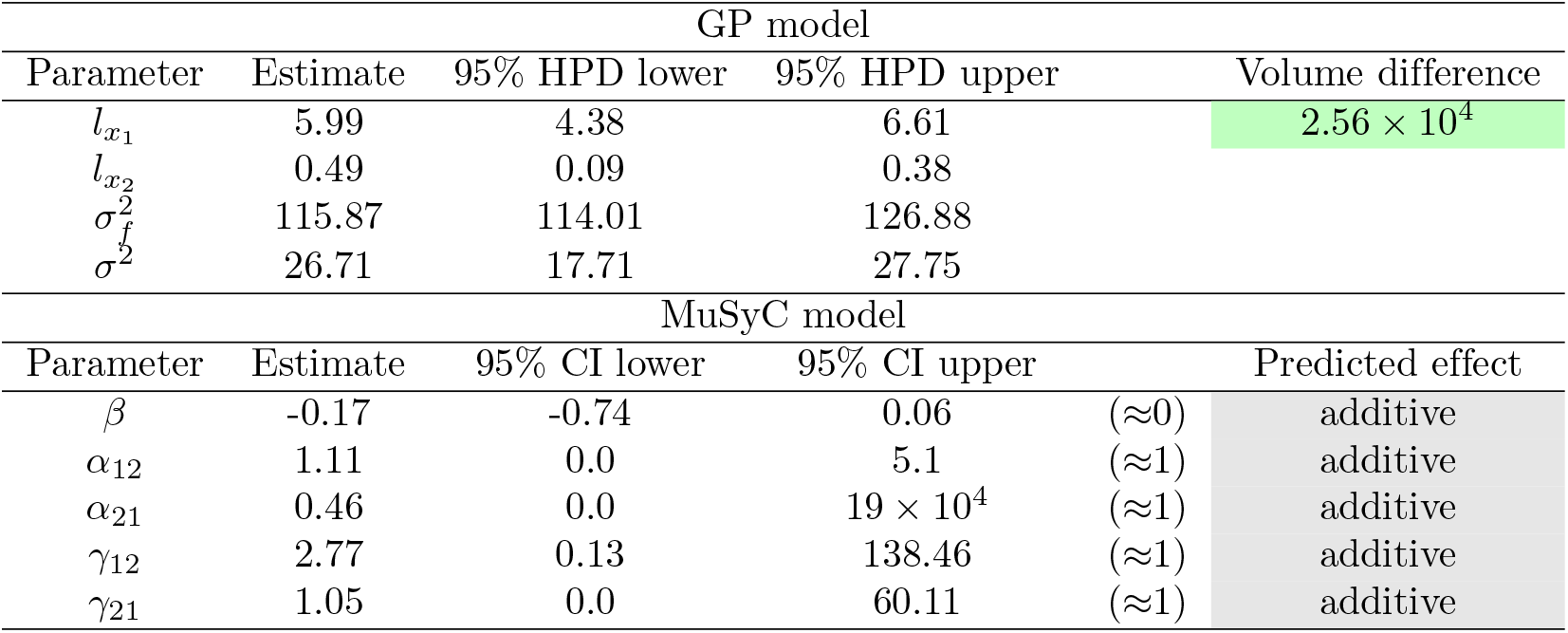
Parameters of the GP and MuSyC models for the Greco data. Also reported are highest posterior density (HPD) estimates from Bayesian inference of the GP and confidence intervals (CI) from maximum likelihood estimation of the MuSyC model.

**Table 3.** Parameters of the GP and MuSyC models for Loewe synergy. Also reported are highest posterior density (HPD) estimates from Bayesian inference of the GP and confidence intervals (CI) from maximum likelihood estimation of the MuSyC model.

| GP model |  |  |  |  |
| --- | --- | --- | --- | --- |
| Parameter | Estimate | 95% HPD lower | 95% HPD upper | Volume difference |
| $l_{x_1}$ | 11.54 | 11.33 | 13.48 | $2.06 \times 10^5$ |
| $l_{x_2}$ | 13.36 | 11.35 | 13.5 | |
| $\sigma_f^2$ | 1.71 | 1.18 | 1.78 | |
| $\sigma^2$ | $4.8 \times 10^{-5}$ | $4.3 \times 10^{-5}$ | $6.5 \times 10^{-5}$ | |
| MuSyC model |  |  |  |  |
| Parameter | Estimate | 95% CI lower | 95% CI upper | Predicted effect |
| $\beta$ | -0.06 | -0.07 | -0.03 | (<0) antagonistic |
| $\alpha_{12}$ | 14.56 | 12.29 | 16.71 | (>1) synergistic |
| $\alpha_{21}$ | 14.56 | 12.93 | 17.32 | (>1) synergistic |
| $\gamma_{12}$ | 0.74 | 0.66 | 0.87 | (<1) antagonistic |
| $\gamma_{21}$ | 0.74 | 0.64 | 0.85 | (<1) antagonistic |

In Figure 2, we provide a more detailed analysis of the differences between the Hand-GP and MuSyC models. The third row shows the synergistic effect surfaces as the difference between the first and second row: green (positive) indicates a synergistic effect and red (negative) indicates an antagonistic effect. We see that the Hand-GP model predicts synergy almost everywhere, whereas the MuSyC model predicts both synergy and antagonism. The bottom row shows the residuals, the difference between the data and the models in the top row. The mean squared errors for the whole surface are similar for the GP and MuSyC models, 13.07 and 13.71 respectively. As can be seen from Figure 2g, the residuals are higher for the GP model around zero doses, but the overall landscape of the residuals appears marginally better in the case of GP model.

**Figure 2.**
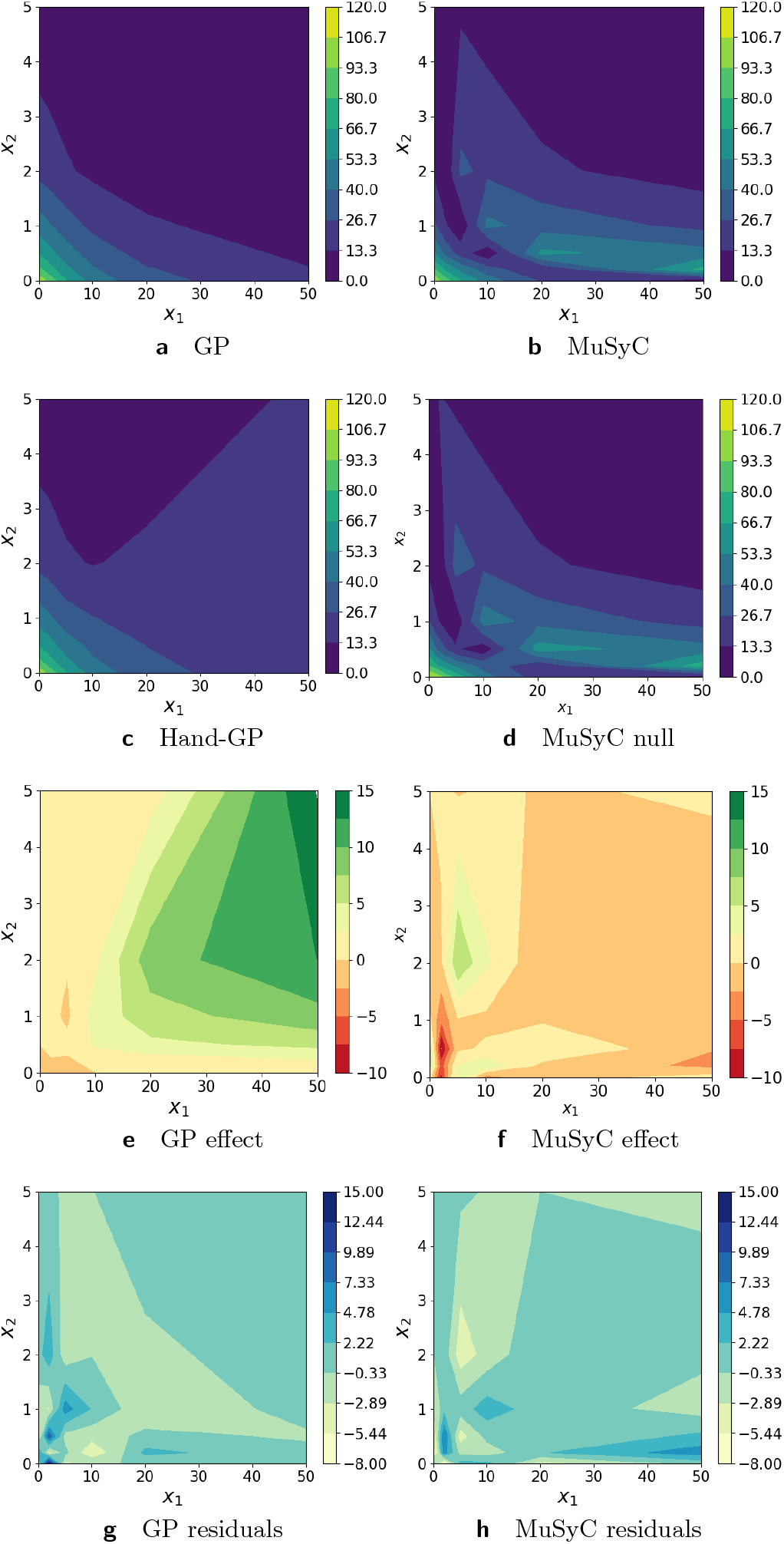
Analysis of Greco simulated data with the Hand-GP (left column) and MuSyC (right column) models. Top row shows the fitted response surfaces. For Hand-GP this is a fit to the non-parametric GP model; for MuSyC a fit to the parametric MuSyC model. The second row shows the null reference models. For Hand-GP this is the Hand construction derived from the fitted monotherapeutic responses from the top row; for MuSyC a fit to a constrained MuSyC model. The third row shows the synergistic effect surfaces as the difference between the first and the second row. The bottom row shows the residuals, the difference between the data and the fits from the top row.

### Simulated data with Loewe synergy and antagonism

As the Greco data set was only mildly synergistic, we generated two data sets with stronger effects (synergistic and antagonistic). Details on the simulated data and parameter values used to generate them can be found in the supplementary material. Figure 3 and Table 3 present results for the synergistic data set. Both the Hand-GP and the MuSyC models predict only synergy as the effect surface is always positive, unlike Figure 2 where the MuSyC model predicted antagonism for some doses. The stronger synergy is reflected in the volume difference of the Hand-GP model which was 2.57 × 10^4^ for the Greco data and now is 2.06 × 10^5^. Curiously, while the effect surfaces in Figure 2 show stronger estimated synergy for the MuSyC model, the synergy parameters in Table 3 indicate antagonism in both efficacy parameter *β* and the Hill slope parameters *γ*_12_ and *γ*_21_.

**Figure 3.**
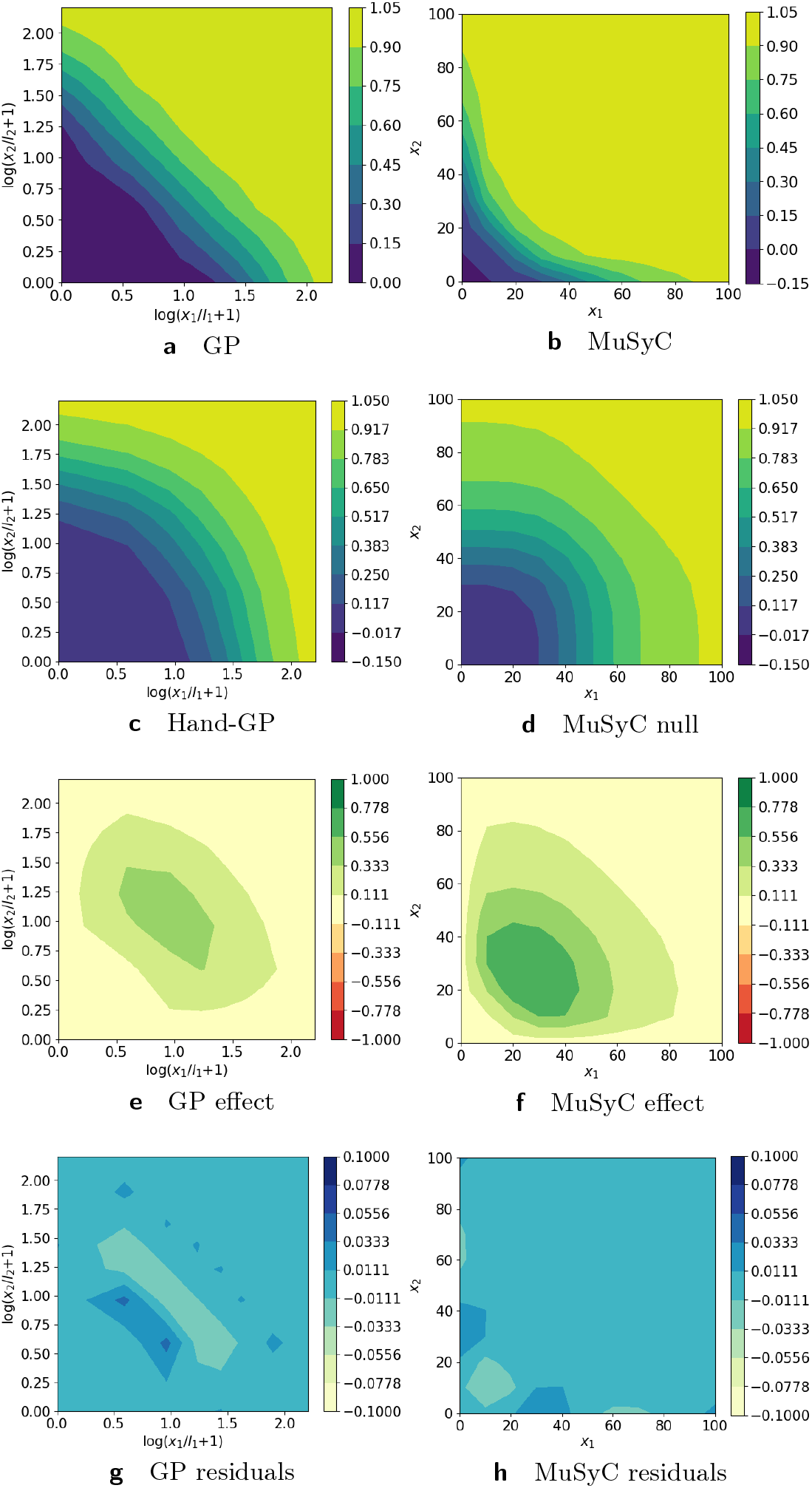
Analysis of Loewe synergy simulated data with Hand-GP (left column) and MuSyC (right column) models. Top row shows the fitted response surfaces. For Hand-GP this is a fit to the non-parametric GP model; for MuSyC a fit to the parametric MuSyC model. The second row shows the null reference models. For Hand-GP this is the Hand construction derived from the fitted monotherapeutic responses from the top row; for MuSyC a fit to a constrained MuSyC model. The third row shows the synergistic effect surfaces as the difference between the first and second row. The bottom row shows the residuals, the difference between the data and the fits from the top row.

Figure 4 and Table 4 present results for the antagonistic data set. Both the Hand-GP and the MuSyC models predict only antagonism as the effect surface is always negative. Concurrently, the synergy parameters of the MuSyC model in Table 3 indicate antagonism in the potency parameters *α*_12_ and *α*_21_ and additivity in the other parameters. Overall, both models perform very well for this data set.

**Figure 4.**
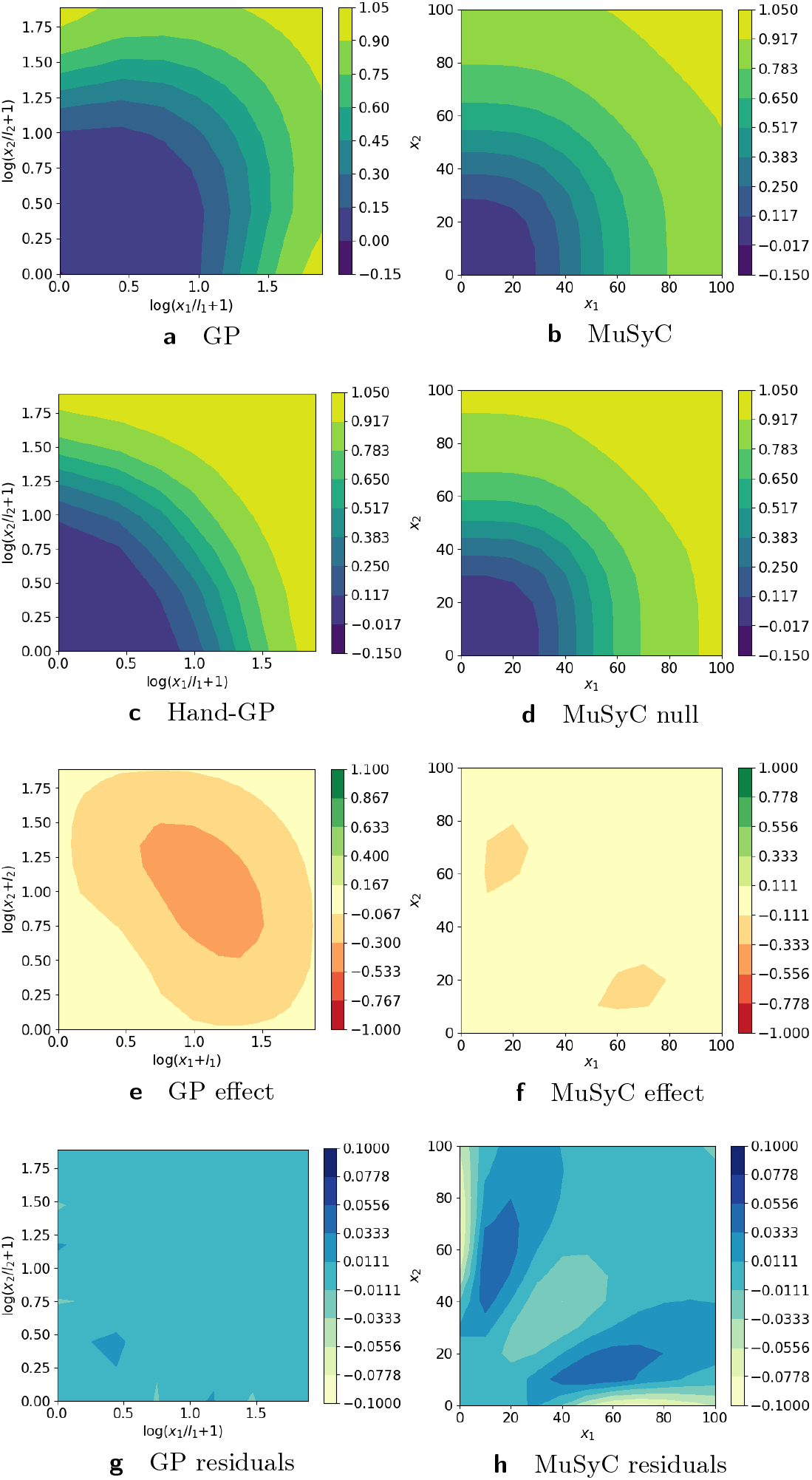
Analysis of Loewe antagonism simulated data set with with Hand-GP (left column) and MuSyC (right column) models. Top row shows the fitted response surfaces, for Hand-GP this is a fit to the non-parametric GP model; for MuSyC a fit to the parametric MuSyC model. The second row shows the null reference models. For Hand-GP this is the Hand construction derived from the fitted monotherapeutic responses from the top row; for MuSyC a fit to a constrained MuSyC model. The third row shows the synergistic effect surfaces as the difference between the first and second row. The bottom row shows the residuals, the difference between the data and the fits from the top row.

**Table 4.**
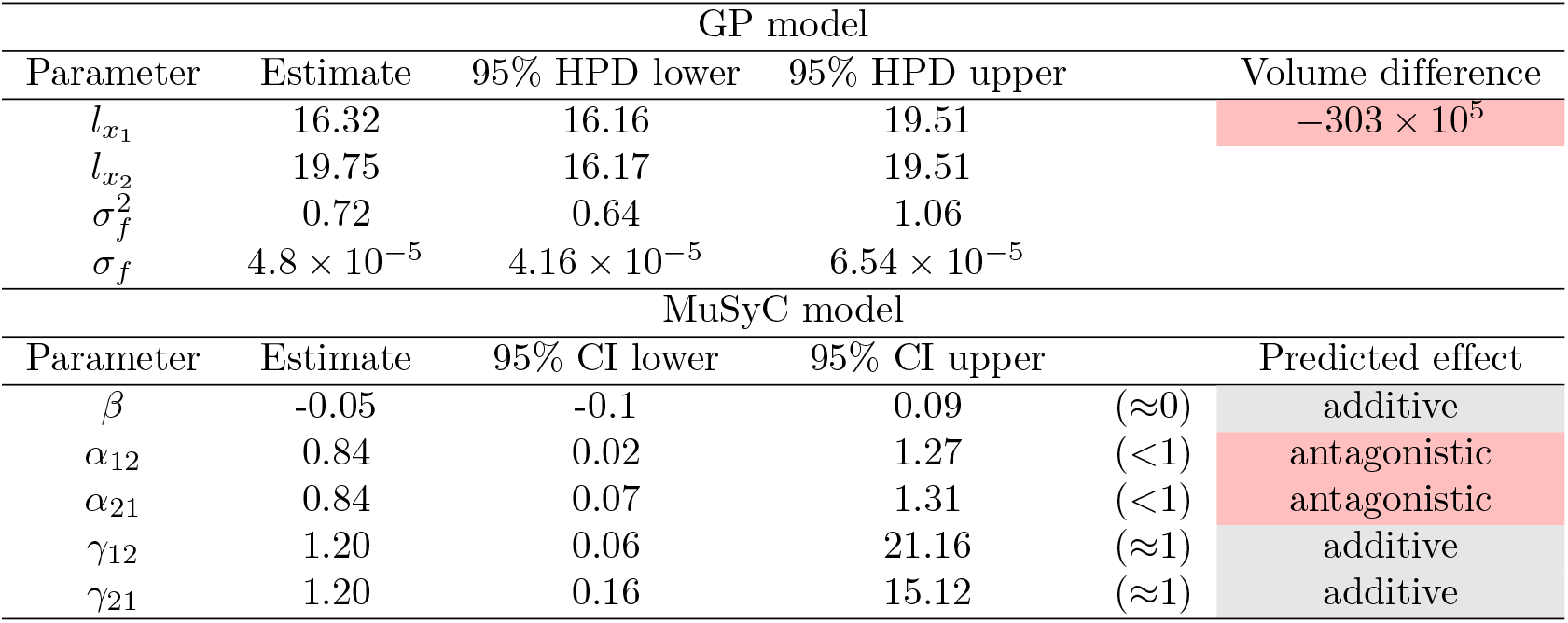
Parameters of GP and MuSyC models for Loewe antagonism. Also reported are highest posterior density (HPD) estimates from Bayesian inference of the GP and confidence intervals (CI) from maximum likelihood estimation of the MuSyC model.

### Experimental data from Chou and Talalay

The data were originally published by Yonetani and Theorell in 1964 [23] and re-analyzed in Chou and Talalay in 1984 [4]. The data are examples of mutually exclusive and non-exclusive inhibitors. The mutually exclusive data contain the inhibition of horse liver alcohol dehydrogenase by the inhibitors ADP-ribose and ADP and the mutually non-exclusive by the inhibitors *o*-phenanthroline and ADP. In our analysis we compare the Hand-GP model to the MuSyC model and the reproduced combination index from the Median Effect model by Chou and Talalay 1984 [4, Figure 3, 5]. We follow the analysis of the Median Effect model, and show predictions from all models for the diagonal rays as indicated in [3, Tab. 1, 2]. However, while the Median Effect model is only fit to the monotherapeutic and diagonal ray data, the Hand-GP and MuSyC models are fit to all 36 dose-response data points.

**Figure 5.**
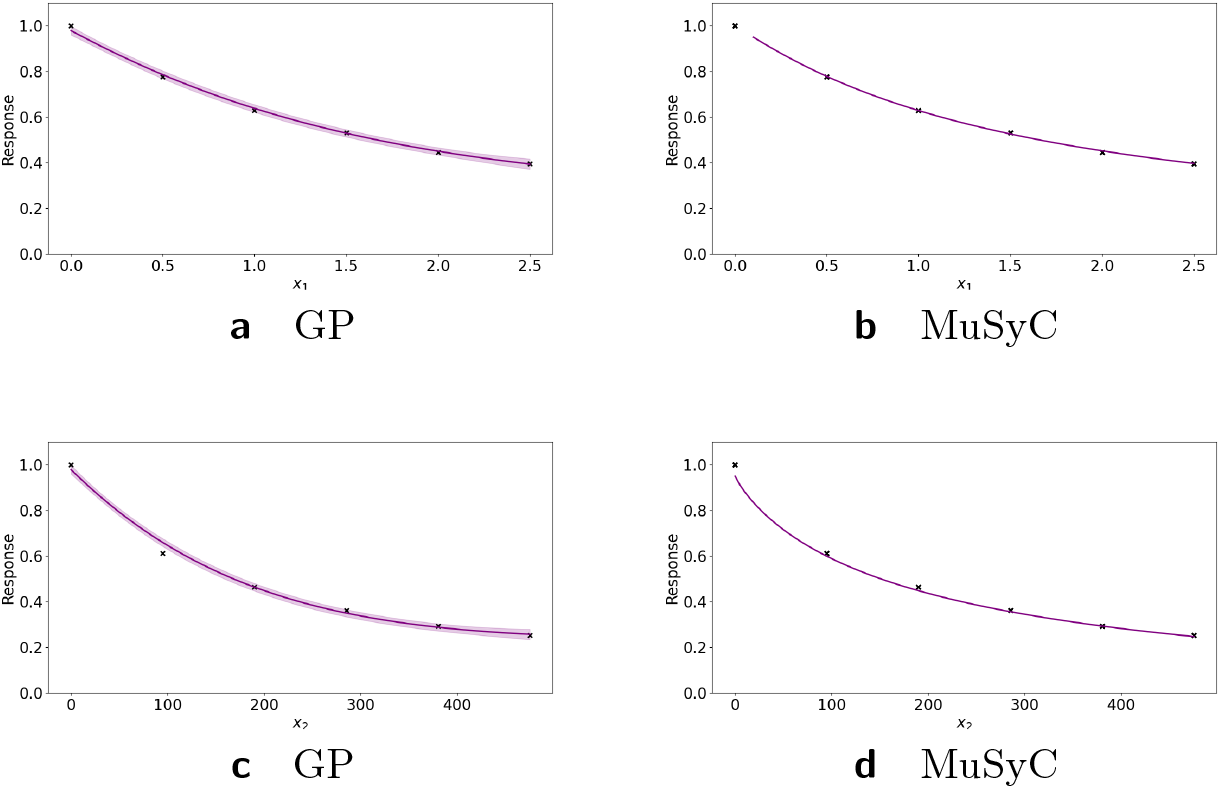
Analysis of mutually exclusive inhibitors data set with with Hand-GP (left column) and MuSyC (right column) models. The figure illustrates monotherapeutic slices from the estimated surfaces. The response is the fractional inhibition of horse liver alcohol dehydrogenase, *x*_1_ is the concentration ADP-ribose and *x*_2_ the concentration ADP.

From Figure 6(a) and (c) we can see that the Hand-GP model predicts antagonism at low doses of both drugs and synergy which at high doses of both drugs (along the diagonal ray). In Figure 6 (e) we present the combination index reproduced from [4] which shows the switch from antagonism to synergy as well, although the combination index is mostly close to 1 which indicates additivity. The MuSyC model predicts mostly antagonistic effects which are close to additivity with a large uncertainty in the parameter *β* and an unidentified upper bound for the parameter *γ*_12_ which predicts synergy. Such uncertainty can be due to the large number of parameters (12) of the MuSyC model relative to the number of data points (36). The results from the Hand-GP model agree with the results from [23]: at lower doses indicating antagonism and at higher doses indicating synergy. All of the models indicate that the overall effect is close to zero (or combination index close to 1 for the Median Effect model) which indicate additivity of the inhibitory effects of ADP-ribose and ADP. Note the different scales of the Hand-GP and MuSyC model in comparison to the scale of the combination index reproduced from [4].

**Figure 6.**
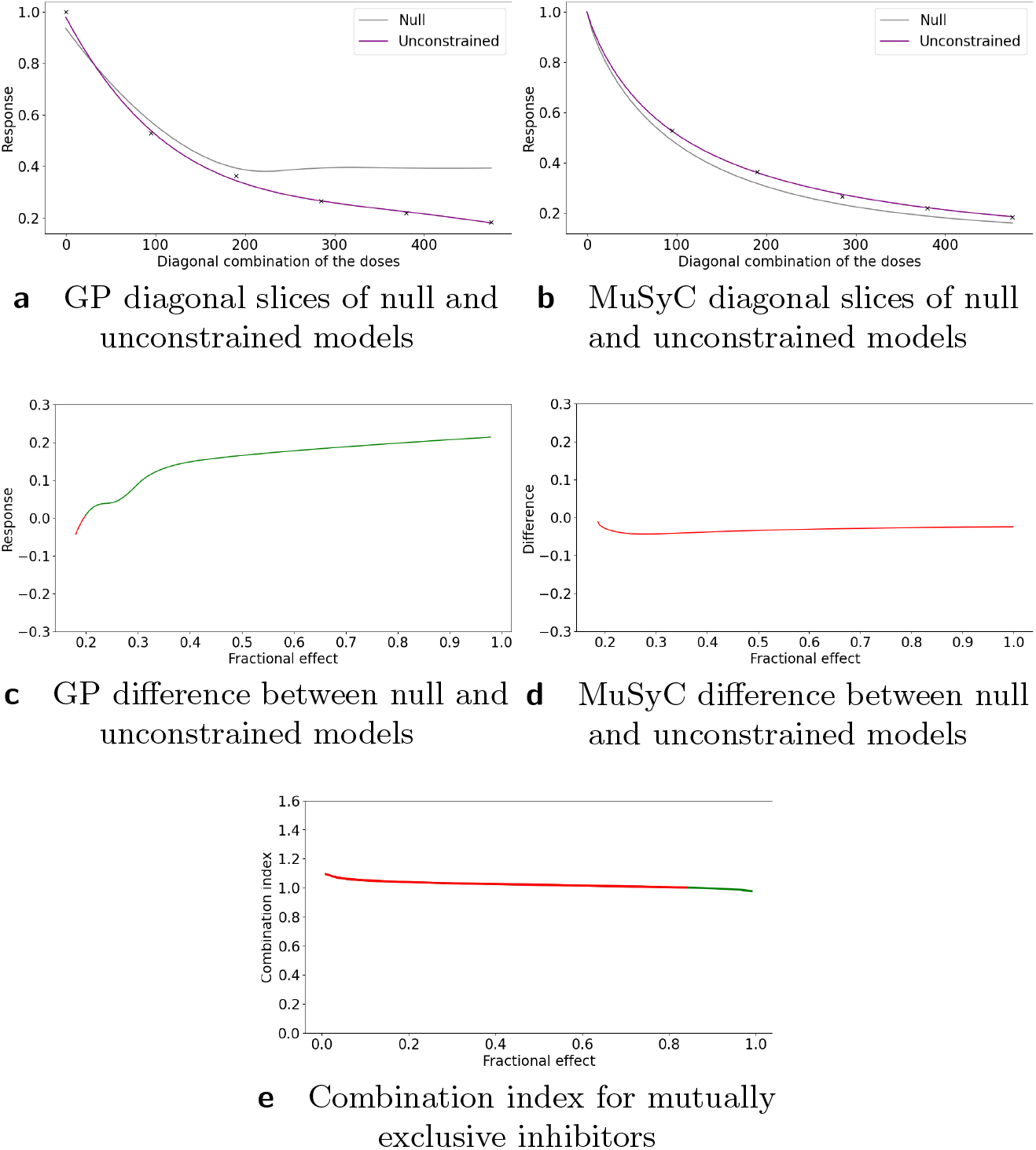
Analysis of mutually exclusive inhibitors data with Hand-GP (left column) and MuSyC (right column) models. Estimated effect with the original Median Effect model analysis is shown in the bottom plot. Top row shows the fitted slices of the surfaces along the diagonal. For Hand-GP this is a fit to the non-parametric GP model; for MuSyC a fit to the parametric MuSyC model. Corresponding null models are in grey and unconstrained models in purple. The middle row shows the difference between the unconstrained and null models. Estimated synergy is shown in green and antagonism in red. Bottom panel shows the combination index reproduced from [4].

**Table 5.** Parameters of GP and MuSyC models for mutually exclusive inhibitors. Also reported are highest posterior density (HPD) estimates from Bayesian inference of the GP and confidence intervals (CI) from maximum likelihood estimation of the MuSyC model.

| GP model |  |  |  |  |  |
| --- | --- | --- | --- | --- | --- |
| Parameter | Estimate | 95% HPD lower | 95% HPD upper |  | Volume difference |
| $l_{x_1}$ | 1.7 | 1.42 | 2.28 | | 3159.9 |
| $l_{x_2}$ | 471.73 | 466.65 | 478.89 | | |
| $\sigma_f^2$ | 1.01 | 0.98 | 1.02 | | |
| $\sigma$ | $2.7 \times 10^{-4}$ | $9.49 \times 10^{-5}$ | $2.1 \times 10^{-3}$ | | |
| MuSyC model |  |  |  |  |  |
| Parameter | Estimate | 95% CI lower | 95% CI upper |  | Predicted effect |
| $\beta$ | $-1.37 \times 10^4$ | $-1.0 \times 10^5$ | $3.8 \times 10^4$ | (<0) | antagonistic |
| $\alpha_{12}$ | 0.01 | 0.0 | 0.01 | (<1) | antagonistic |
| $\alpha_{21}$ | 0.0 | 0.0 | 0.45 | (<1) | antagonistic |
| $\gamma_{12}$ | 18.66 | 18.66 | inf | (>1) | synergistic |
| $\gamma_{21}$ | 0.76 | 0.01 | 47.37 | (<1) | antagonistic |

### Inhibition of alcohol dehydrogenase by two mutually non-exclusive inhibitors

The second study in [23] concerned the inhibition of horse liver alcohol dehydrogenase by two competitive, mutually non-exclusive inhibitors: *o*-phenanthroline and ADP and was re-analyzed in [4] using the Median Effect model. In Figure 8(e) we present combination index reproduced from [4]. We re-analyze these data using the Hand-GP and MuSyC models. Using the Hand-GP model we obtain antagonism for the combination of low doses and synergism for the combination of high doses which agrees with the analysis in [4]. The MuSyC model predicts synergy for the parameters *α*_12_ and *α*_21_ (change of effective dose) and antagonism for *γ*_12_ and *γ*_21_, change of the Hill coefficient. Figure 8(d) illustrates that along the diagonal MuSyC model predicts mostly synergy. Complete surface analysis for both models can be found in the supplementary material. In Figure 7 we can see that the MuSyC model does not fit the monotherapeutic part of the data well, while the GP fits look good. Both models fit the diagonal of the surface quite well, which is illustrated in Figure 8. In this case the results of the Hand-GP model agree with the analysis in [4].

**Figure 7.**
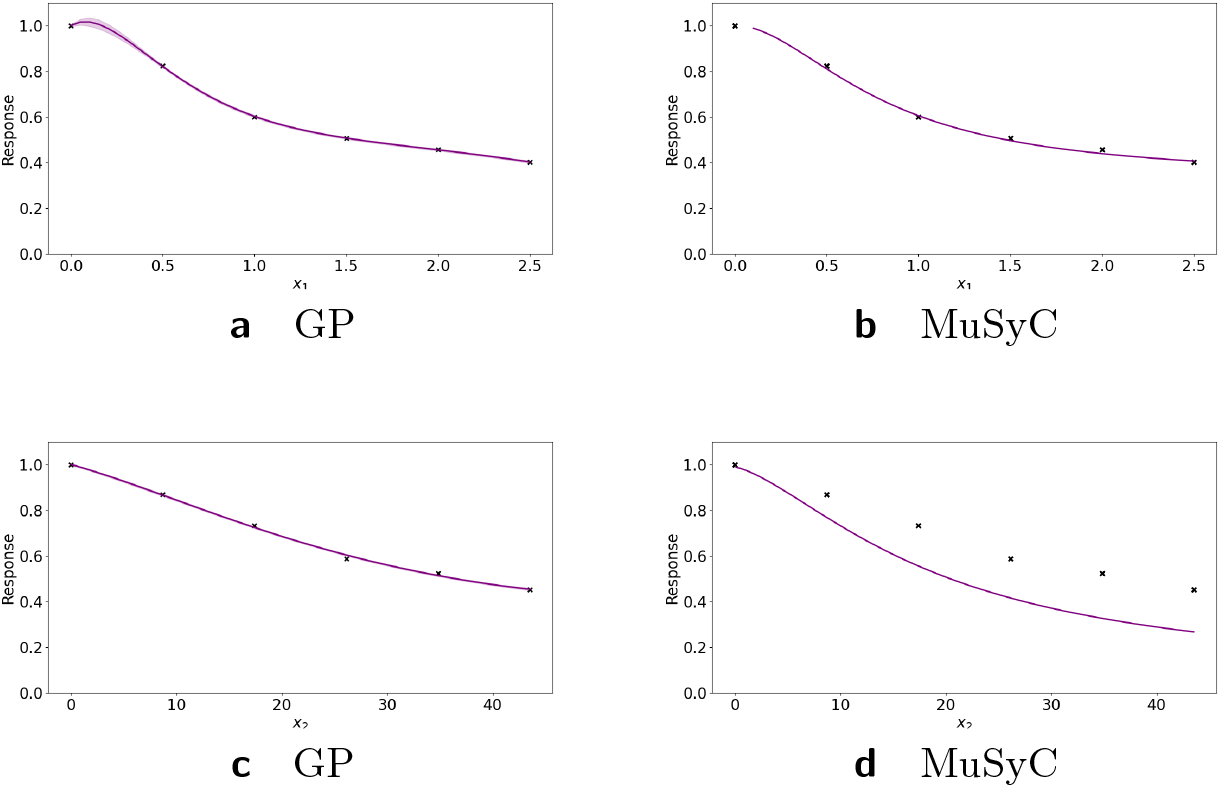
Analysis of mutually non-exclusive inhibitors data set with with Hand-GP (left column) and MuSyC (right column) models. The figure illustrates monotherapeutic slices from the estimated surfaces. The response is the fractional inhibition of horse liver alcohol dehydrogenase, *x*_1_ is the concentration *o*-phenanthroline and *x*_2_ the concentration ADP.

**Figure 8.**
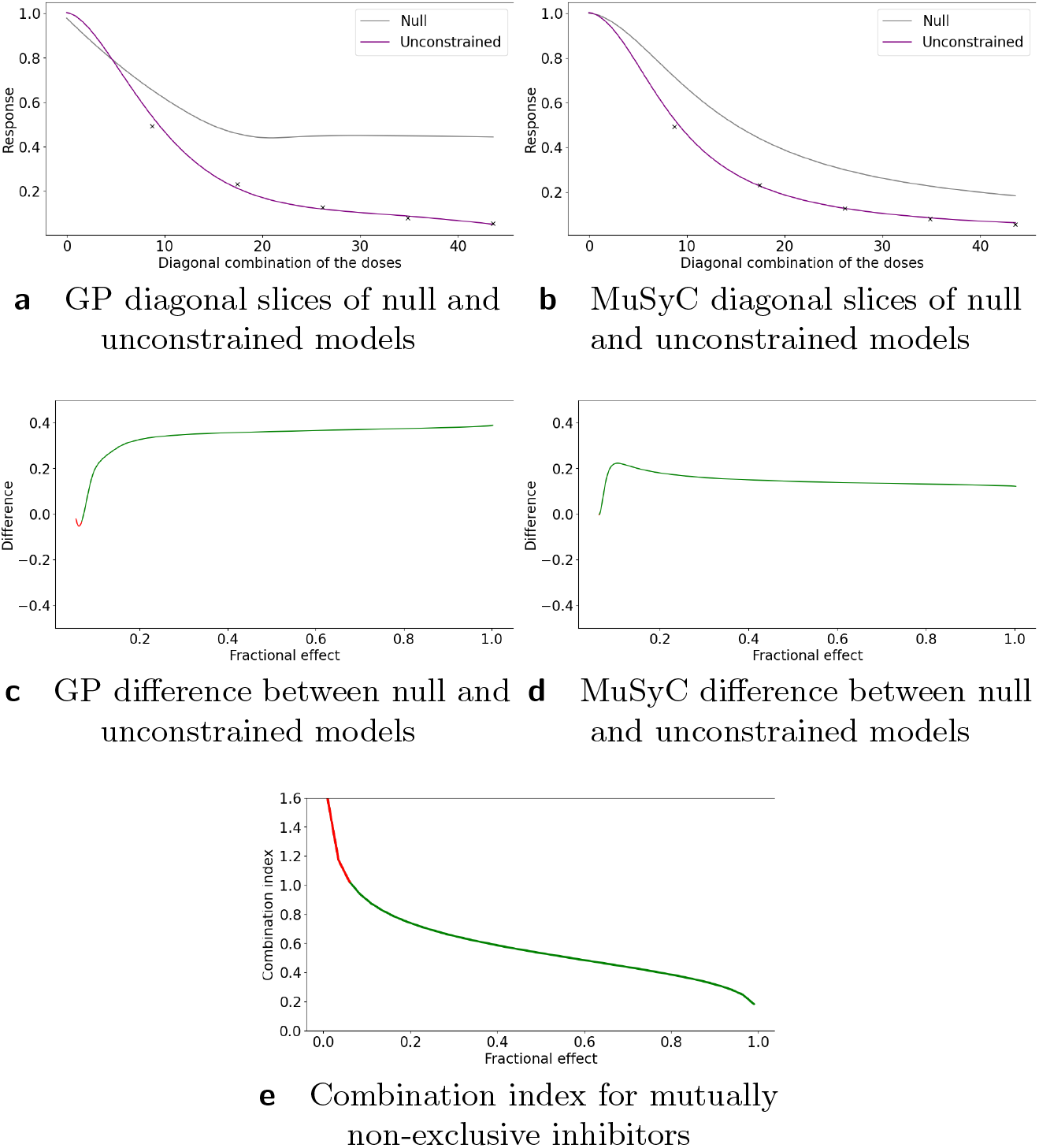
Analysis of mutually non-exclusive inhibitors data with Hand-GP (left column) and MuSyC (right column) models. Estimated effect with the original Median Effect model analysis are shown at the bottom plot. Top row shows the fitted slices of the surfaces along the diagonal. For Hand-GP this is a fit to the non-parametric GP model; for MuSyC a fit to the parametric MuSyC model. Corresponding null models are in grey and unconstrained models in purple. The middle row shows the difference between predicted effects by unconstrained and null models. Estimated synergy effect is shown in green and antagonism in red. Bottom panel shows the combination index reproduced from [4].

## Discussion

We introduced the Hand-GP model, a combination of the Hand principle for constructing null reference models with a Gaussian process using a new logarithmic squared exponential kernel. We demonstrated performance of the Hand-GP model on multiple benchmark data sets, both simulated and experimental ones. We compared the Hand-GP model with the recent parametric MuSyC model and with the well-established Loewe, Bliss and Median Effect models. In all cases the Hand-GP model performed very well and we obtained correct or expected predictions of synergistic effects. Our comparison of the proposed method to the MuSyC model shows that parametric models- although useful and interpretable - cannot account for some dose-response surfaces that are observed in practice. If the experimental data deviates too much from the assumed parametric curve/surface then the predicted effect can be incorrect. In such situations a non-parametric approach can be of help to predict an interaction effect.

**Table 6.**
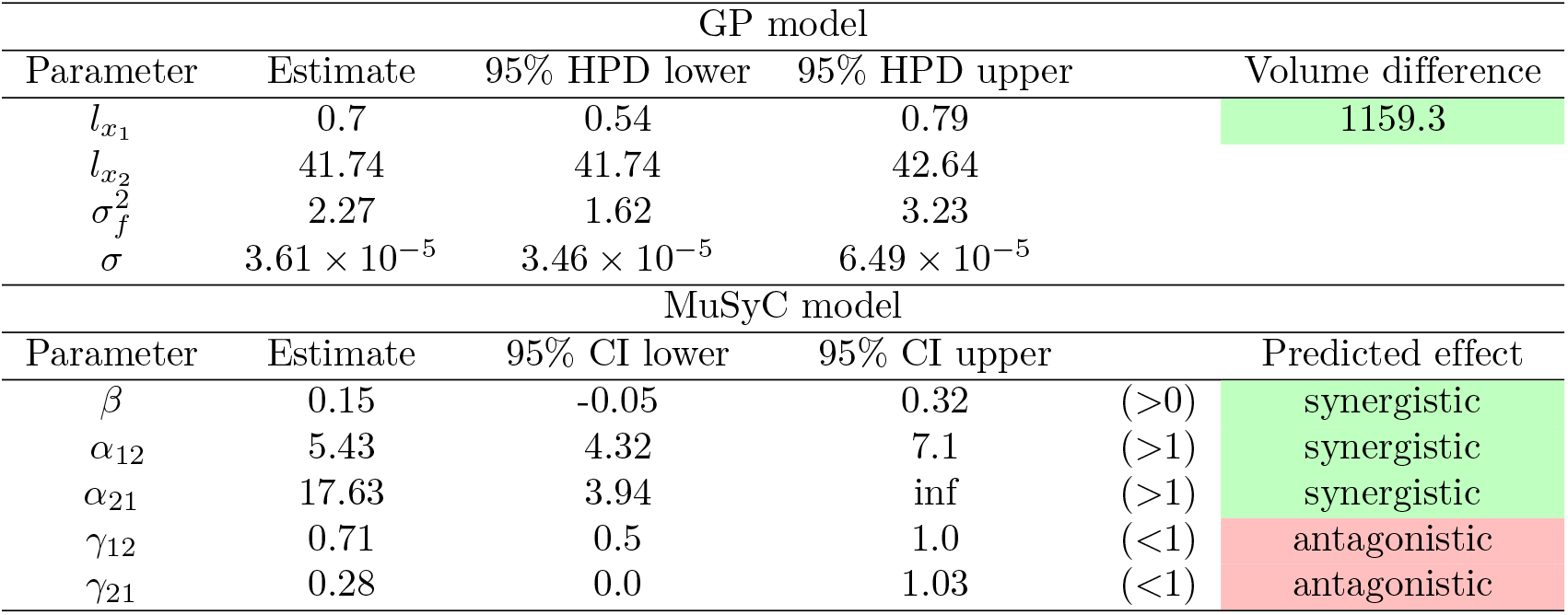
Parameters of GP and MuSyC models for mutually non-exclusive inhibitors. Also reported are highest posterior density (HPD) estimates from Bayesian inference of the GP and confidence intervals (CI) from maximum likelihood estimation of the MuSyC model.

| GP model |  |  |  |  |
| --- | --- | --- | --- | --- |
| Parameter | Estimate | 95% HPD lower | 95% HPD upper | Volume difference |
| $l_{x_1}$ | 0.7 | 0.54 | 0.79 | 1159.3 |
| $l_{x_2}$ | 41.74 | 41.74 | 42.64 | |
| $\sigma_f^2$ | 2.27 | 1.62 | 3.23 | |
| $\sigma$ | $3.61 \times 10^{-5}$ | $3.46 \times 10^{-5}$ | $6.49 \times 10^{-5}$ | |
| MuSyC model |  |  |  |  |
| Parameter | Estimate | 95% CI lower | 95% CI upper | Predicted effect |
| $\beta$ | 0.15 | -0.05 | 0.32 | (>0) synergistic |
| $\alpha_{12}$ | 5.43 | 4.32 | 7.1 | (>1) synergistic |
| $\alpha_{21}$ | 17.63 | 3.94 | inf | (>1) synergistic |
| $\gamma_{12}$ | 0.71 | 0.5 | 1.0 | (<1) antagonistic |
| $\gamma_{21}$ | 0.28 | 0.0 | 1.03 | (<1) antagonistic |

While we showcased the Hand-GP model for binary and time-fixed compound interactions, the model can be easily extended to include more than two compounds or time as an additional component in the Gaussian process. Especially time has received only limited attention in the synergy literature, but see [10]. More generally, viewing time as part of the phenotype, there can be many different types of changes in phenotype in response to perturbations, e.g. changes of cell shape or spatial rearrangements of the cell. All these changes can be flexibly modeled within the GP framework, leveraging research in the machine learning community [1, 2]. Further extensions are possible to incorporate various experimental scenarios. For example, when the data are too noisy and thus the observations are non-monotonic, a constrained Gaussian processes framework can be used [12, 15, 16] such that monotonicity is enforced in the GP. Ultimately, we view our model proposal as a building block in a pipeline that would predict the phenotype in a cell type in response to multiple perturbations. The flexibility of Gaussian processes make such a vision more likely than parametric modeling approaches.

## Conclusion

In this paper, we proposed a non-parametric approach to dose-response synergy modeling based on Gaussian processes and the Hand reference model. We proposed a new kernel function that operates on the log-scale of the doses and takes into account the specifics of the cellular response to perturbations that often depend on the logarithm of the dose. We estimate not only monotherapeutic dose-response curves, but rather fit a complete dose-response surface to the data and use the resulting robust estimates of the monotherapeutic dose-response curves to construct a null reference response surface. The model can be extended to incorporate more inputs, such as time or location and is flexible enough to function as a building block in a pipeline that predicts cellular response to multiple perturbations.

## Method

### Gaussian process dose-response model

In this section, we construct Gaussian process (GP) models for dose-response data. The models are based on Gaussian processes regression with multi-dimensional input. For ease of understanding we start with the univariate model. Let *x* be a dose and *y* the response. We assume that the dose-response relationship follows

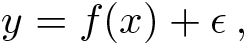

where *ϵ* is a noise term and *f*(*x*) is represented by a GP: *f*(*x*) ~ *GP* (0, *k*(*x, x′*)) with *k*(·, ·) a kernel function. A common choice of kernel for smooth functions is the squared exponential:

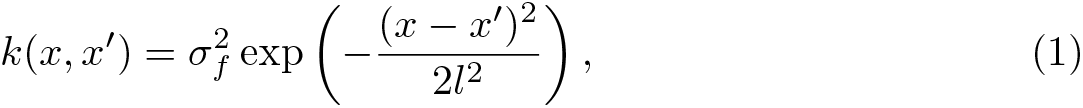

where 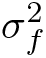 and *l*, correspondingly the variance and the length scale, are hyperparameters of the kernel. The variance determines how far from its mean the GP function deviates on average, and the length scale determines the smoothness of the function. The squared exponential kernel is stationary, and defines an infinitely differentiable function. [18] show that this kernel is universal in the sense that under some conditions, this kernel can learn any continuous structure given enough data [6]. The disadvantage of this kernel for dose-response modeling is that usually responses are affected by the logarithm of the dose, and a simple log transformation of dose is not desirable since we would need to take the logarithm of zero as dose zero is typically included in an experiment. We solve this problem by proposing the following kernel:

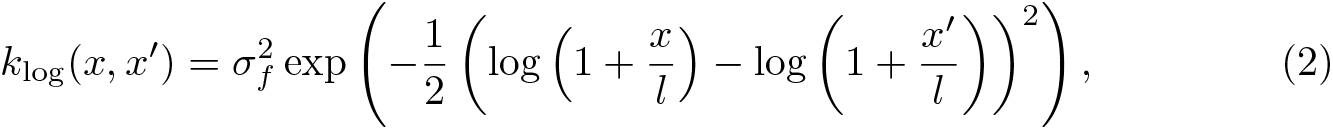

where *l* still has the interpretation of a length scale and 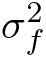 the amplitude of the GP function. Equation (2) defines a kernel for individual dose-response functions, i.e., for input *x* ∈ ℝ, *f*: ℝ → ℝ. For doses {*x, x′*} ≪ *l*, we have log(1 + *x/l*) ≈ *x/l*, and the kernel (2) is indistinguishable from the squared exponential kernel (1). Figure 9 shows samples from the squared-exponential and logarithmic kernels.

**Figure 9.**
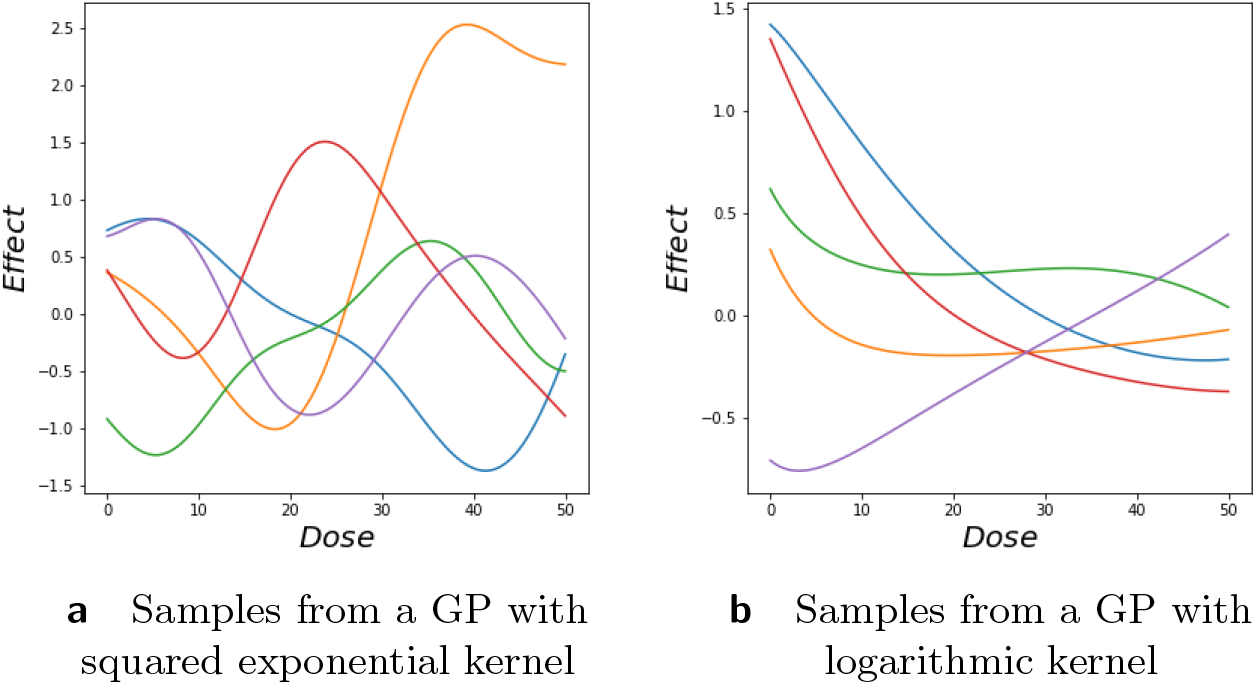
Samples from logarithmic kernel. (a) Samples from the logarithmic squared exponential kernel with *l* = 10.0 and 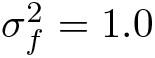. (b) Samples from the squared exponential kernel with *l* = 10 and 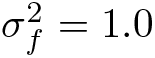.

The generalization to bivariate input is straightforward. Let ***x*** = (*x*_1_, *x*_2_) ∈ ℝ^2^, *f*: ℝ^2^ → ℝ, then *f*(***x***) = *GP*(0, *k*_log_(***x***, ***x′***)), where

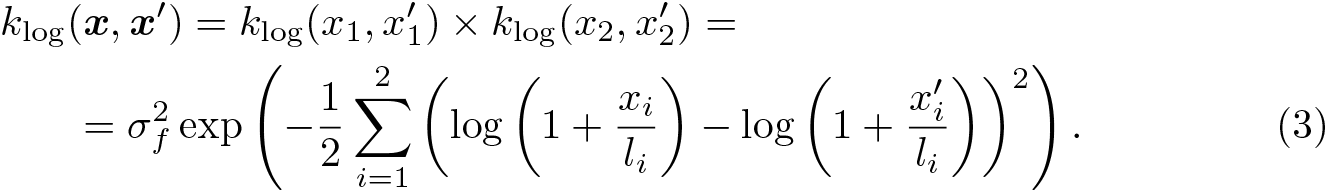

The kernel in Equation (3) is constructed by multiplying two logarithmic kernels. This kernel structure intuitively means that *f*(*x*_1_, *x*_2_) is only expected to be similar to some other function value 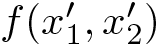 if *x*_1_ is close to 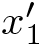 and *x*_2_ is close to 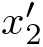. Note that in Equation (3) each dimension has its own length-scale parameter. In the dose-response framework the ratio of these length scales *l*_1_*/l*_2_ corresponds to the potency ratio, i.e., the ratio with which two drugs can substitute each other. The complete observational model with noise now reads

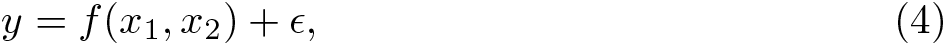

where 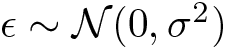 with hyperparameter *σ*^2^ the noise strength. In all, the bivariate GP model has four hyperparameters ***θ*** = {*σ*, *σ*_*f*_, *l*_1_, *l*_2_}.

Let us denote by ***y*** = (*y*_1_,… , *y*_*n*_) the observations *y*_*i*_ corresponding to the dose combinations ***x***_*i*_ = (*x*_1*i*_, *x*_2*i*_), which are combined into matrix *X*. Let 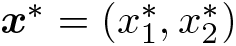 be a test point with corresponding prediction *f*\*. Following [19], we have

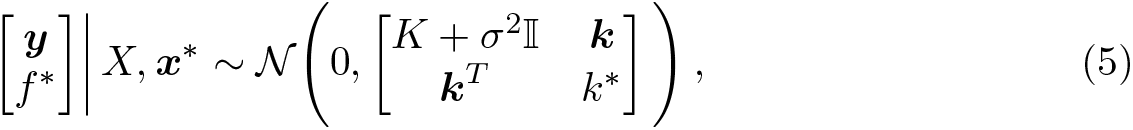

where *K* is the *n* × *n* kernel matrix with elements *K*_*ij*_ = *k*_log_(***x***_*i*_, ***x****_j_*), ***k*** an *n* × 1 vector with elements *k*_*i*_ = *k*_log_(***x***_*i*_, ***x***\*), and *k*\* = *k*_log_(***x***\*, ***x***\*). This leads to the posterior distribution

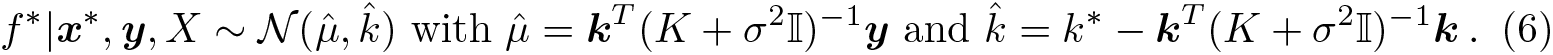

Given hyperparameters and a data set with responses for various dose combinations, we can use Equation 6 to compute the posterior mean response and credible intervals for any possible dose combination ***x***\*.

We used a Bayesian approach to find the posterior distributions of the hyperparameters by sampling the hyperparameters using Hamiltonian Monte Carlo (HMC) methods [5]. A HMC sampling algorithm uses gradients of the target distribution, which allows for much more efficient exploration of the parameter space than MCMC algorithms relying on random walk proposals. We used the TensorFlow Probability library for the implementation of HMC. We ran the algorithm with 1000 burn-in steps and 100000 samples. Target acceptance rate was set to 75%. Note that this acceptance rate is high for standard MCMC algorithms with random walk proposals such as Metropolis-Hastings, but common for MCMC algorithms that use Hamiltonian dynamics. We found that 5 leapfrog steps worked well in all of the applications considered in this paper. The proposal step size was tuned individually and chosen from the set {0.01, 0.03, 0.05, 0.07, 0.1, 0.2, 0.3, 05, 0.7, 1.0}. In each case the largest step that achieved the target acceptance rate was chosen. We used weakly informative priors for the hyperparameters of the kernel. We used a Gamma(*α,β*) distribution for the hyperparameters of the kernel, length scale, and variance. For the length scale we chose *α* and *β* such that *α/β* is equal to the maximal dose of a drug scaled by a constant *c* and *α/β*^2^ is 0.1× half of the maximal dose of the drug scaled by a constant *c*. Constant *c* reflects whether the effect changes in the observed data points. One strategy to set *c* is to set it to the ratio between maximal and minimal effect reached on the monotherapeutic data. For the variance we choose *α* and *β* such that *α/β* is equal to half of the maximum effect reached and *α/β*^2^ is equal to 0.1×half of the maximum effect reached. A Gamma(0.14, 1.14) prior was used for the noise variance in all applications except Greco data. For Greco data we used Gamma(2, 1) prior. This difference is due to the different scale of the data. Examples of the samples obtained with HMC can be found in supplementary information. As one can see, the chains are well mixed and converged. The uncertainty about the parameter values is based on the 95% highest posterior density intervals obtained from the HMC samples.

### Null reference model from the Hand principle

We are interested in whether two drugs are *synergetic* or *antagonistic*, i.e. whether the mixture has stronger/weaker effect compared to what we would have expected if the drugs had no interaction. Null reference models are designed such that there is no interaction between drugs. There are three desirable properties that null reference models should satisfy [20]:

- *Sham combination principle:* a drug does not interact with itself. Hence the combination of a drug with itself leads to neither synergy nor antagonism;
- *Commutativity:* swap of drugs should not change the results;
- *Associative property:* combining combination drugs should be the same as directly combining drugs at corresponding ratios.

Sinzger et al. 2019 [20] compare various popular null reference models and show that the Hand model is biochemically the most plausible as it is the only model satisfying all three desirable properties. The Hand model construction is close to that of Loewe, which is detailed in Lederer et al. 2018. The Hand model can be viewed as an infinitesimal version of the Loewe model, thereby solving the commutativity issues that plague models based on Loewe additivity. In Algorithm 1 we present the implementation of the Hand-GP model and in Figure 10 we provide an illustration of the Hand construction. Note that *f*^−1^(·) has to be approximated numerically since it is impossible to find the inverse of a Gaussian process analytically. However, if we use a large enough number of test points for predicting a Gaussian process, we get a very good numerical approximation. For Algorithm 1 we need to choose *N*_1_, the number of partitions of the dose *x*_1_, and *N*_2_, the number of partitions of the dose *x*_2_. We choose *N*_1_ and *N*_2_ depending on the number of test points for the Gaussian process model and where the applied doses *x*_1_ and *x*_2_ are located among these test points. By the test points here we mean the points in which we made prediction for the Gaussian process regression, denoted as ***x***\*. For choosing *N*_1_ we find the closest point among the test points to *x*_1_ and choose *N*_1_ as the number of test points we have between 0 dose and closest test point to *x*_1_. *N*_2_ is chosen in the same way considering dose *x*_2_.

**Figure 10.**
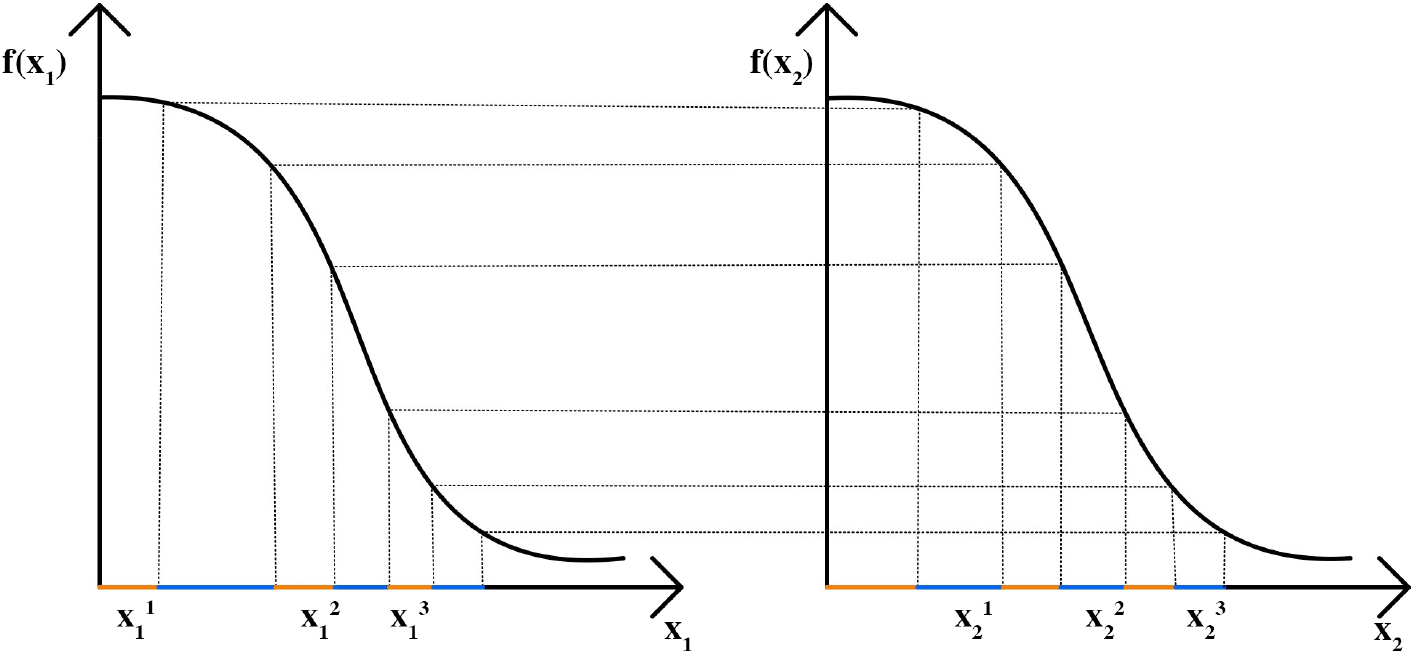
Illustration of the Hand model. In this case doses *x*_1_ and *x*_2_ are split into *N*_1_ = *N*_2_ = 3 parts and the partitions are applied sequentially.

**Algorithm 1.**
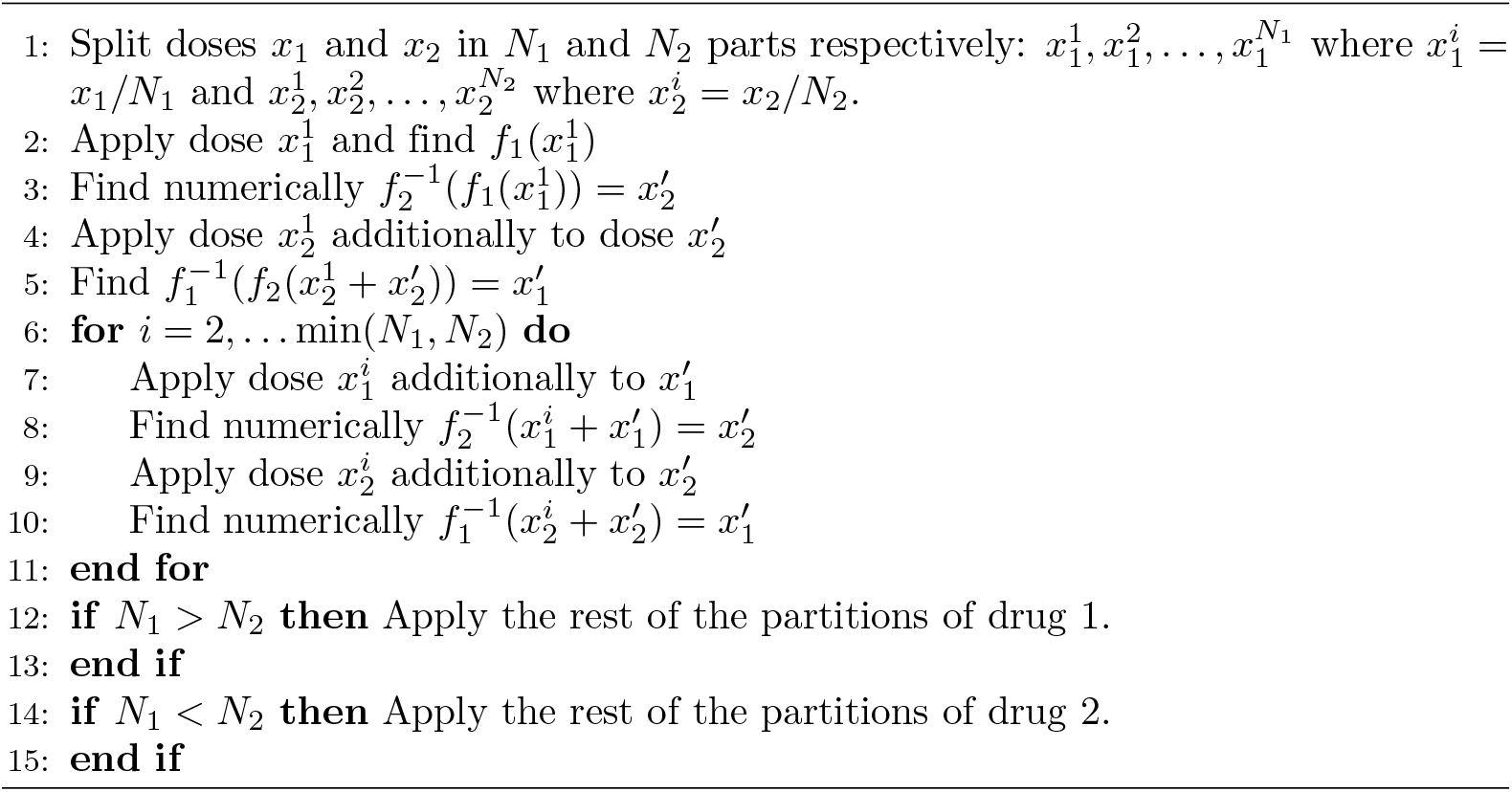
Implementation of Hand GP model

## Availability of data and materials

The datasets analyzed in this paper and the source code are available at https://github.com/YuliyaShapovalova/HandGP. Archived version: 10.5281/zenodo.4623762.

## Author’s contributions

All authors contributed to the design of the study. YS implemented the Hand-GP model and performed the data analysis. TD implemented the Median Effect model. All authors wrote or contributed to the writing of the manuscript. All authors read and approved the final manuscript.

## A Supplementary figures

**Figure 11.**
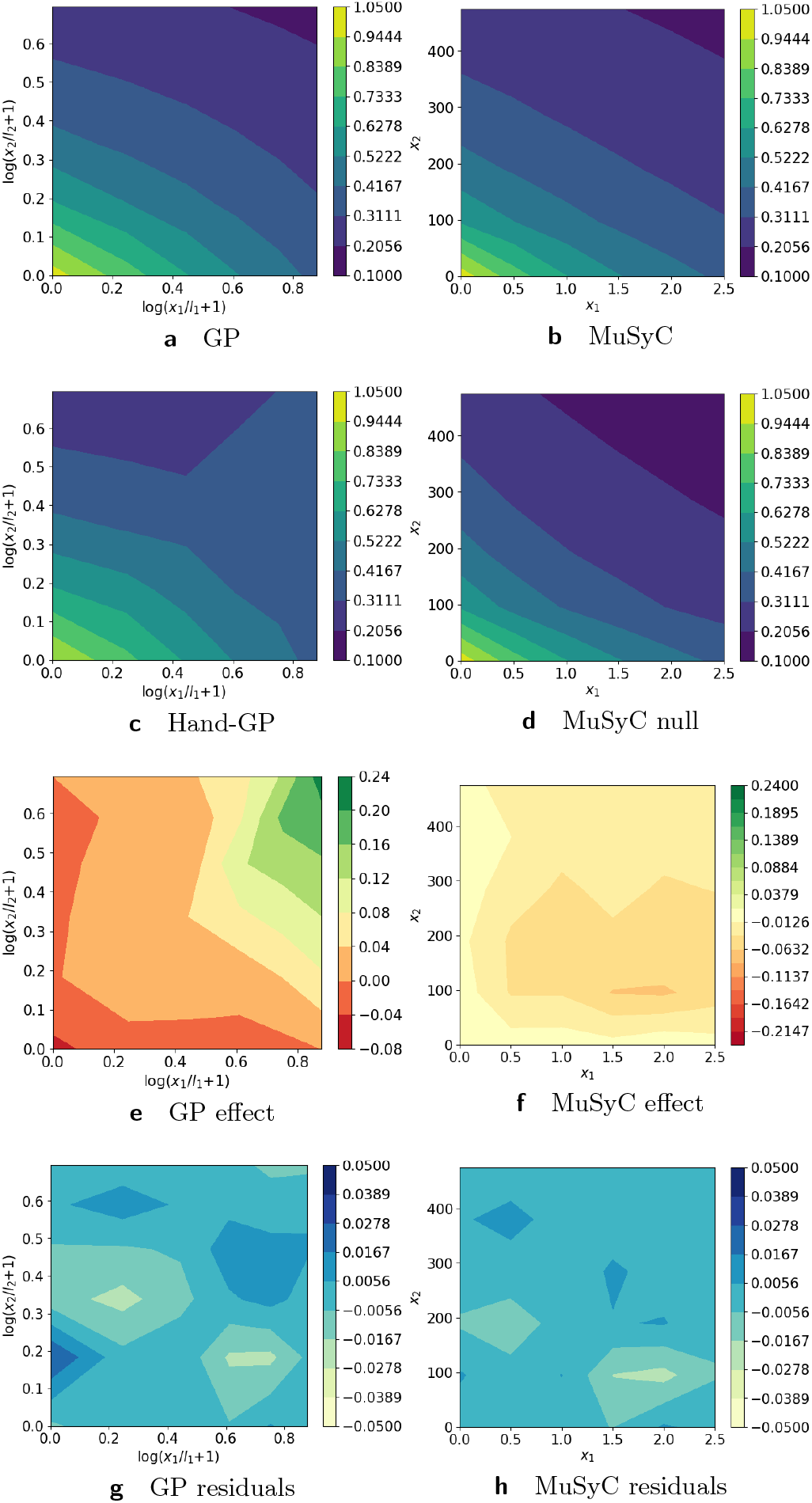
Analysis of mutually exclusive inhibitors data [23] set with with Hand-GP (left column) and MuSyC (right column) models. Top row shows the fitted response surfaces, for Hand-GP this is a fit to the non-parametric GP model for MuSyC a fit to the parametric MuSyC model. The second row shows the null reference models. For Hand-GP this is the Hand construction derived from the fitted monotherapeutic responses from the top row. For the MuSyC model this is a fit to a constrained MuSyC model. The third row shows the synergistic effect surfaces as difference between the first and second row. The bottom row shows the residuals, the difference between the data and the fits from the top row.

**Figure 12.**
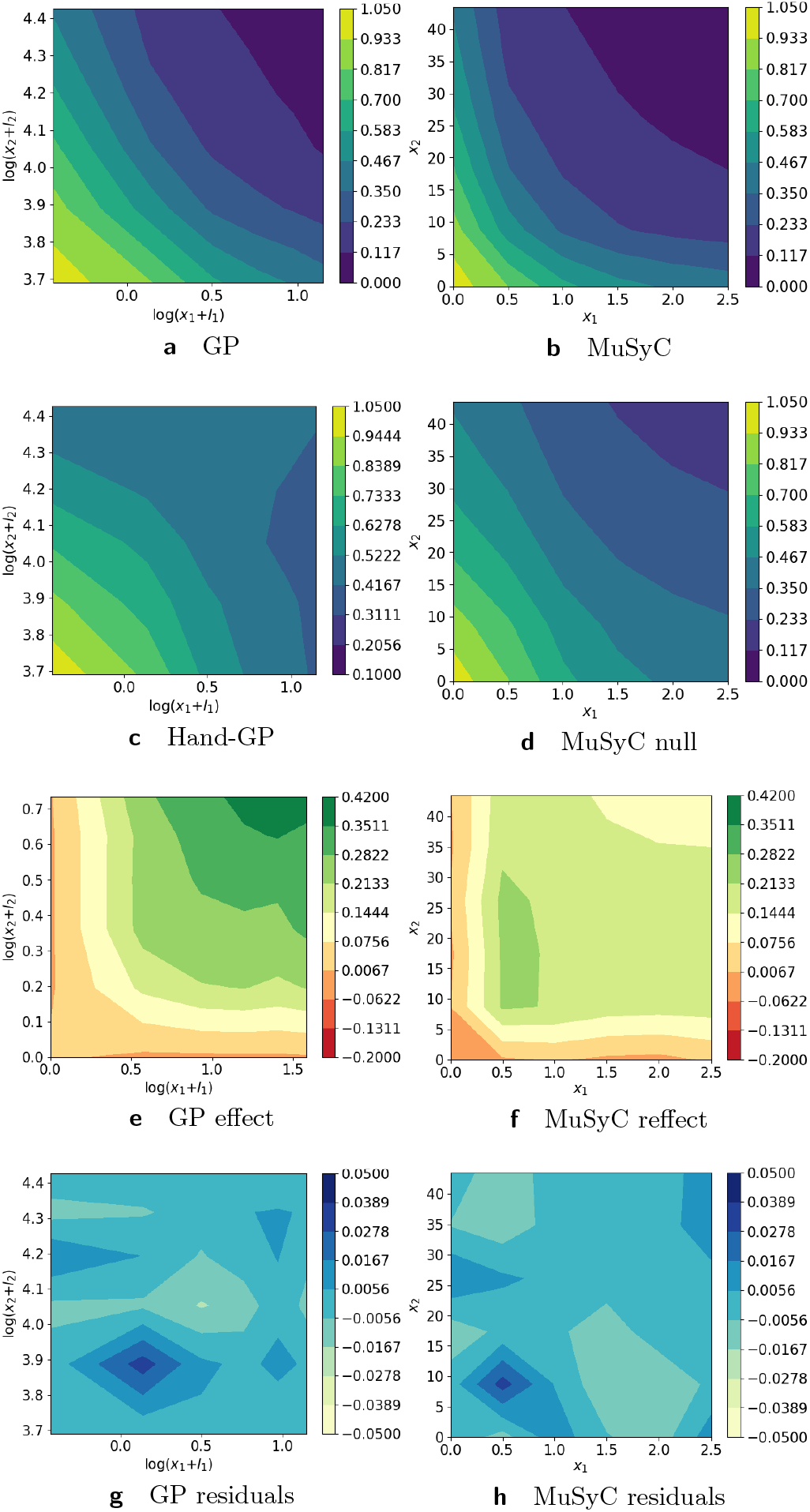
Analysis of mutually non-exclusive inhibitors data [23] set with with Hand-GP (left column) and MuSyC (right column) models. Top row shows the fitted response surfaces, for Hand-GP this is a fit to the non-parametric GP model for MuSyC a fit to the parametric MuSyC model. The second row shows the null reference models. For Hand-GP this is the Hand construction derived from the fitted monotherapeutic responses from the top row. For the MuSyC model this is a fit to a constrained MuSyC model. The third row shows the synergistic effect surfaces as difference between the first and second row. The bottom row shows the residuals, the difference between the data and the fits from the top row.

**Figure 13.**
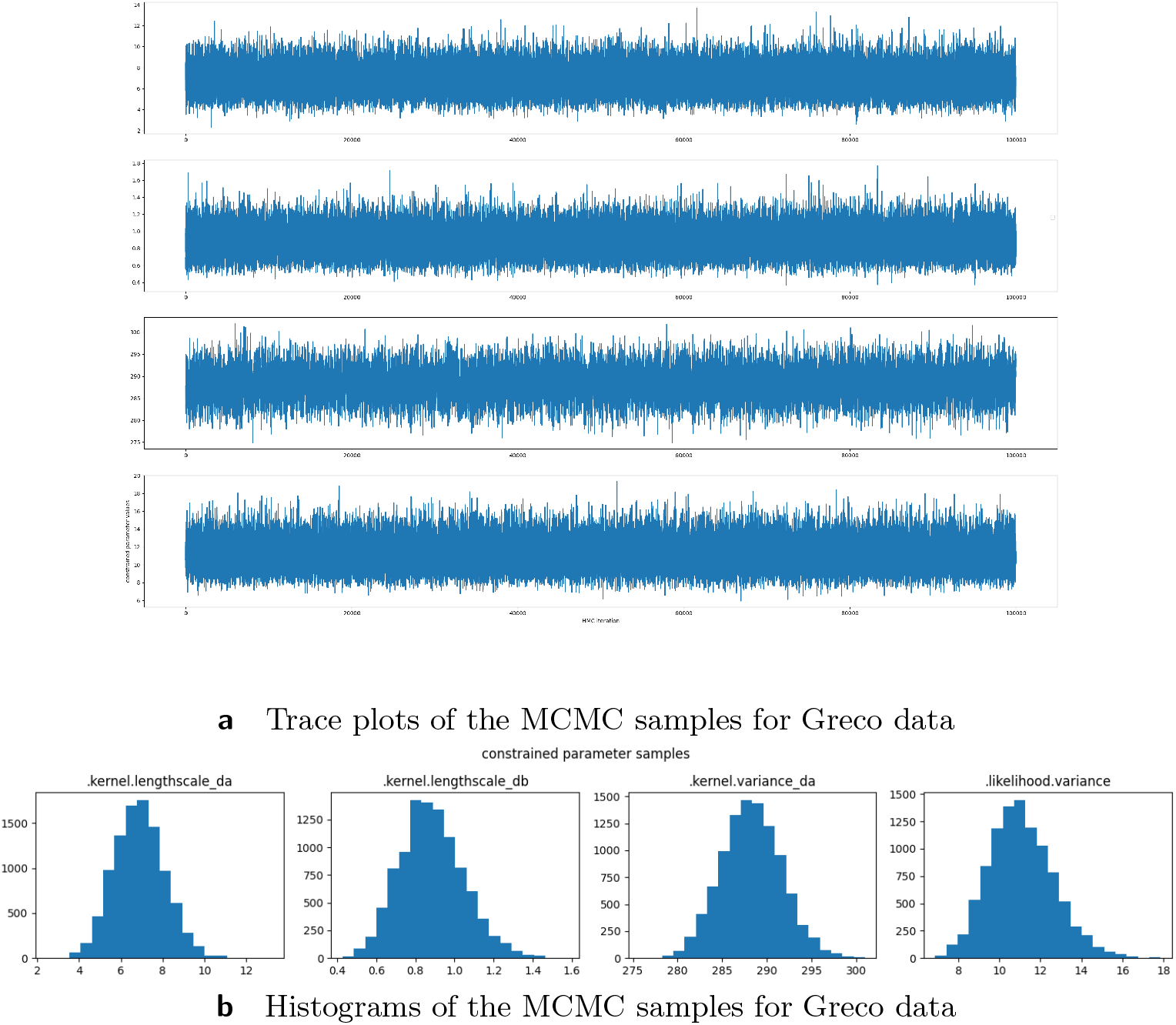
MCMC results for Greco data set. The samples are obtained using Hamiltonian Monte Carlo with leap frog step 5, step size 0.7, target acceptance rate 0.75 and adaptation rate 0.1. We used HMC sampler from TensorFlow Probability. Since the number of the hyperparameters is small, it is not hard to get good performance of HMC in terms of convergence and mixing.

## B Simulated data Loewe synergy and Loewe antagonism

We use approach of [21] to generate bidirectional synergistic and antagonistic data set in Loewe additivity frameworks. We use the code that provided with the paper to generate these data [21]. The parameters for generating the data: 1) synergy *Emax*_1_ = 1, *EC*50_1_ = 50, *H*_1_ = 4, *Emax*_2_ = 1, *EC*50_2_ = 50, *H*_2_ = 4, *Int*_12_ = −.9, *Int*_21_ = −.9, *EC*50*Int*_12_ = 25, *EC*50*Int*_21_ = 25; 2) antagonism *Emax*_1_ = 1, *EC*50_1_ = 50, *H*_1_ = 4, *Emax*_2_ = 1, *EC*50_2_ = 50, *H*_2_ = 4, *Int*_12_ = 1, *Int*_21_ = 1, *EC*50*Int*_12_ = 25, *EC*50*Int*_21_ = 25. No noise was added to these data.

